# Mechanism of IRSp53 inhibition by 14-3-3

**DOI:** 10.1101/430827

**Authors:** David J. Kast, Roberto Dominguez

## Abstract

Filopodia are precursors of dendritic spines and polarized cell migration. The I-BAR-domain protein IRSp53 is an essential regulator of filopodia dynamics that couples Rho-GTPase signaling to cytoskeleton and membrane remodeling, playing essential roles in neuronal development and cell motility. Here, we describe a mechanism whereby phosphorylation-dependent inhibition of IRSp53 by 14-3-3 counters membrane binding and activation by Cdc42 or downstream cytoskeletal effectors. Phosphoproteomics, quantitative binding studies and crystal structures show that 14-3-3 binds to two pairs of phosphorylation sites in IRSp53. Using bicistronic expression we obtained a heterodimer of IRSp53 in which only one subunit is phosphorylated, and show that each subunit of the IRSp53 dimer independently binds a 14-3-3 dimer. A FRET-sensor assay developed using natively phosphorylated and 14-3-3-binding competent IRSp53 purified from mammalian cells reveals opposite conformational changes in IRSp53 upon binding of activatory (Cdc42, Eps8) *vs*. inhibitory (14-3-3) inputs.

## Introduction

A tight correlation between plasma membrane and actin cytoskeleton dynamics is a common feature of many cellular functions, including cell migration, organelle trafficking and endo/exocytosis. Bin/Amphiphysin/Rvs (BAR) domain proteins are essential spatio-temporal coordinators of signaling events to actin cytoskeleton and membrane dynamics. The superfamily of BAR domain protein comprises a diverse group of multi-functional effectors, containing N- and/or C-terminal to the membrane-binding BAR domain additional signaling and protein-protein or protein-membrane interaction modules, including SH3, PX, PH, RhoGEF and RhoGAP domains ^1,2^. Depending on the shape of the BAR domain, three subfamilies are distinguished: the classical crescent-shaped BAR, the more extended and less curved F-BAR and the inverse-curvature I-BAR subfamilies. The most prominent member of the I-BAR subfamily is IRSp53 (also known as BAIAP2), a protein implicated in the formation of plasma membrane protrusions such as filopodia ^3,4,5,6,7,8,9,10^ and dendritic spines ^11, 12, 13, 14, 15, 16^. IRSp53 functions under the control of Cdc42 ^4,6,9,10,17,18^, and recruits to the plasma membrane a long list of cytoskeletal effectors and synaptic scaffolds, including Eps8 ^6,10^, Ena/VASP ^4,9,20^, N-WASP ^18^, WAVE ^3,21, 22^, mDia ^8, 23^, PSD-Q5 ^11, 12, 24, 25^ and Shank3 ^11, 17^. As a consequence, IRSp53 is both a key player in normal developmental processes, such as dendritic spine development^16^, myoblast fusion ^26^ and eye development^27^, as well as a frequent factor in diseases, including tumorigenesis ^19, 28^ and neurological disorders ranging from autism spectrum disorder ^29^ and schizophrenia ^30^ to attention deficit hyperactivity disorder ^31^.

IRSp53 features an N-terminal I-BAR domain (residues 1-231) implicated in membrane binding ^32,33, 34^ The I-BAR domain is immediately followed by a partial CRIB motif, which is interrupted by a proline-rich sequence and is thus referred to as the CRIB-PR domain (residues 260–291) ^10^. C-terminal to the CRIB-PR domain, and connected by an 85-amino acid linker rich in serine, threonine and proline residues, IRSp53 presents an SH3 domain (residues 375–437). The SH3 domain mediates most of the interactions of IRSp53 with downstream cytoskeleton effectors. In the inactive state, the SH3 domain binds intramolecularly to the CRIB-PR domain, resulting in a compact, closed conformation ^10^. A transition toward an active, open conformation can be triggered by the binding of either Cdc42 to the CRIB-PR domain or cytoskeletal effectors to the SH3 domain ^10^. Cdc42-dependent activation of IRSp53 in cells results in increased formation of membrane ruffles and filopodia-like structures ^4,6,9,10,17,18^. The region between the CRIB-PR and SH3 domains hosts multiple phosphorylation sites, and some of these sites have been implicated in 14-3-3 binding ^35, 36^, which can lead to loss of cell polarity ^35^. Here, we show that phosphorylation of two pairs of sites within this region triggers the binding of 14-3-3 to IRSp53, which inhibits membrane binding and the interactions of Cdc42 with the CRIB-PR domain and cytoskeletal effectors with the SH3 domain.

### Purification of phosphorylated and 14-3-3-binding-competent pIRSp53

Mammals express seven 14-3-3 isoforms, denoted β, γ, ε, ζ, η, σ, and θ (also called τ), and most isoforms are enriched in brain tissue ^37^, where IRSp53 is also enriched ^7, 16^. IRSp53 has been shown to interact *in vitro* and in cells with several 14-3-3 isoforms, including ζ, γ, θ, and σ ^35, 36, 38, 39^. Here, we established conditions that enhance the amount of IRSp53 that binds 14-3-3 in mammalian cells, and developed a protocol for the purification of this protein in large quantities for biochemical and phosphoproteomics studies (**Fig. 1**). A previous report linked IRSp53 phosphorylation at T340 and T360 to the binding of 14-3-3 ^36^. These authors additionally found that IRSp53 coimunoprecipitates less abundantly with 14-3-3 when cells are treated with LiCl, leading them to suggest that a kinase downstream of GSK3β phosphorylates IRSp53 at 14-3-3-binding sites. However, another study found that a Par1b/MARK2 mutant that accepts *N*^6^-benzyl-ATP[^35^*γ*S] as substrate directly phosphorylates IRSp53 at S366, and this site accounted for the largest fraction of 14-3-3 pulling down with IRSp53 ^35^. Par1b/MARK2 is an AMPK-related kinase ^40^, and AMPK has been also shown to directly phosphorylate IRSp53 at S366 using a similar chemical genetics screening approach ^41, 42^. Since AMPK-related kinases are the sole to be conclusively shown to directly phosphorylate IRSp53, we used serum starvation, known to activate AMPK-dependent phosphorylation ^43^, to enhance IRSp53 phosphorylation in HEK293T cells. As anticipated, serum starvation increased the relative amount of endogenous 14-3-3 (all isoforms included) that coimmunoprecipitates with FLAG-IRSp53, and an optimal starvation time of 2 h pre-lysis was established for future experiments (**Fig. 1a**). Consistent with previous findings ^35,41, 42^, these results indicate that AMPK (or a related kinase) phosphorylates IRSp53, and that the AMPK phosphorylation sites(s) are at least in part responsible for the binding of 14-3-3 ^35^.

**Fig. 1.**
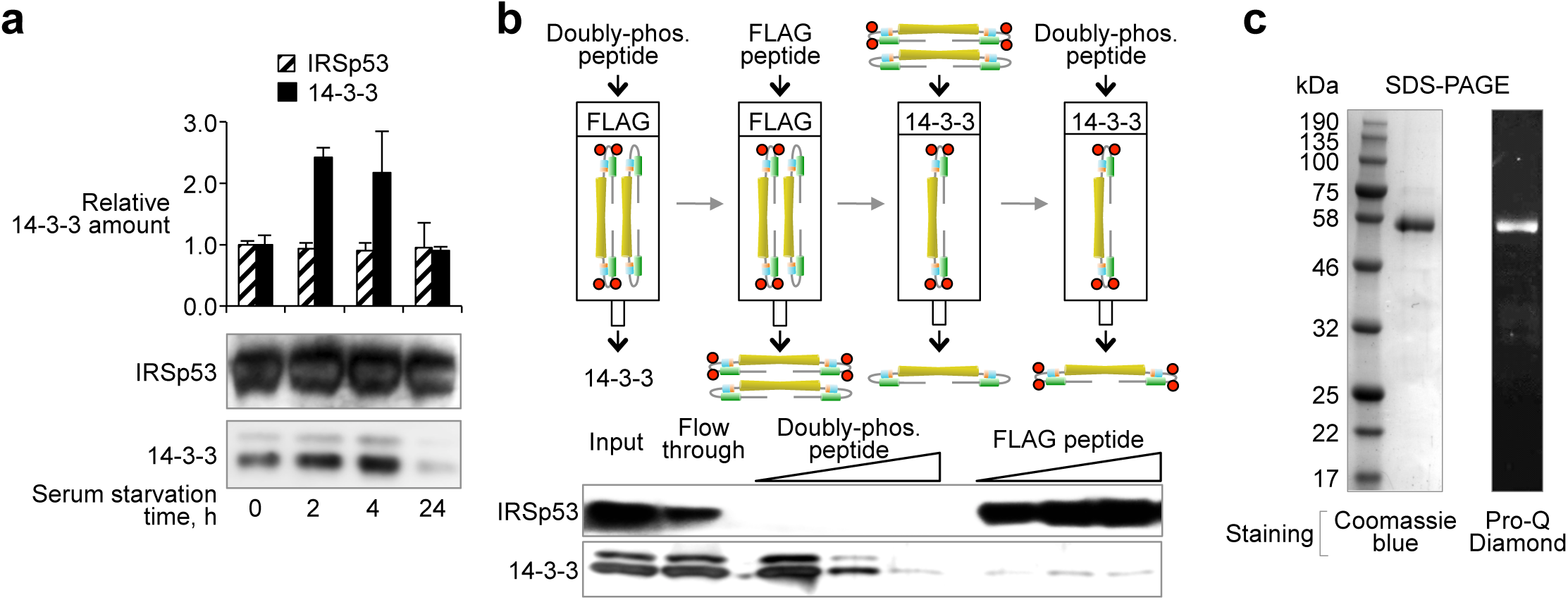
Purification of phosphorylated and 14-3-3-binding-competent pIRSp53. **a** Effect of serum starvation on the relative amount of 14-3-3 (black bars) that coimmunoprecipitates with IRSp53-FLAG (stripped bars) from HEK293T cells. Abundance is reported relative to fed cells expressing IRSp53-FLAG (first column). Error bars are ± s.d. from three independent experiments. Note that serum starvation does not affect the expression of IRSp53-FLAG, but increases the relative amount of 14-3-3 that coimmunoprecipitates with it. An optimal starvation time of 2 h was established from these experiments. **b** Purification of phosphorylated IRSp53-FLAG from HEK293T cells. Cells were serum-starved for 2 h to enhance IRSp53 phosphorylation. Endogenous 14-3-3 bound to IRSp53 is competitively removed by addition of the doubly-phosphorylated IRSp53 peptide Sites 2,3 (**Supplementary Table 1**). The 14-3-3-binding-competent fraction of IRSp53 is then isolated through consecutive affinity purification steps (FLAG-affinity and 14-3-3-affinity), eluting IRSp53 by competition with either FLAG or Sites 2,3 peptides. **c** Coomassie and Pro-Q Diamond-stained SDS-PAGE gels showing that the isolated fraction of IRSp53 is pure and phosphorylated.

We developed a protocol for the purification of phosphorylated and 14-3-3-binding-competent IRSp53 (pIRSp53) from serum-starved HEK293T cells using alternating FLAG- and 14-3-3-affinity steps (**Fig. 1b**). A FLAG peptide and a doubly-phosphorylated IRSp53 synthetic peptide (Sites 2,3 peptide; see below) were used to elute IRSp53 from the affinity columns by competition. The progress of the purification was monitored by Western blot analysis (**Fig. 1b**). SDS-PAGE analysis confirmed the purity of the purified protein, which was additionally shown to be phosphorylated using Pro-Q Diamond gel staining (Molecular Probes, Eugene, OR) (**Fig. 1c**).

### Characterization of the 14-3-3-binding sites of pIRSp53

A survey of phosphoproteomics studies compiled in PhosphoSitePlus ^44^ reveals that IRSp53 is mostly phosphorylated in the flexible, S/T-rich region between the CRIB-PR and SH3 domains (**Fig. 2a**), which seems like an ideal location for the binding of 14-3-3 to inhibit interactions of the CRIB-PR with Cdc42 and the SH3 domain with cytoskeletal effectors. Coincidentally, two studies have identified 14-3-3-binding sites within this region, albeit different from one another: pT340/pT360 ^36^ and S366 and to a much lesser extent a residue within the cluster ^452^SSST^455^ located after the SH3 domain ^35^. We analyzed by mass spectrometry the highly pure, 14-3-3-binding-competent pIRSp53 preparation obtained here (**Fig. 1c**). Five sites were reproducibly phosphorylated in the four samples analyzed (three with 2 h starvation and one without starvation): S325, T340, T360, S366 and a residue within the ^452^SSST^455^ cluster, most likely S454 (**Fig. 2a**, red bars). These sites coincide with those found with the highest frequency in phosphopoteomic studies, and include the sites previously implicated in 14-3-3 binding ^35, 36^.

**Fig. 2.**
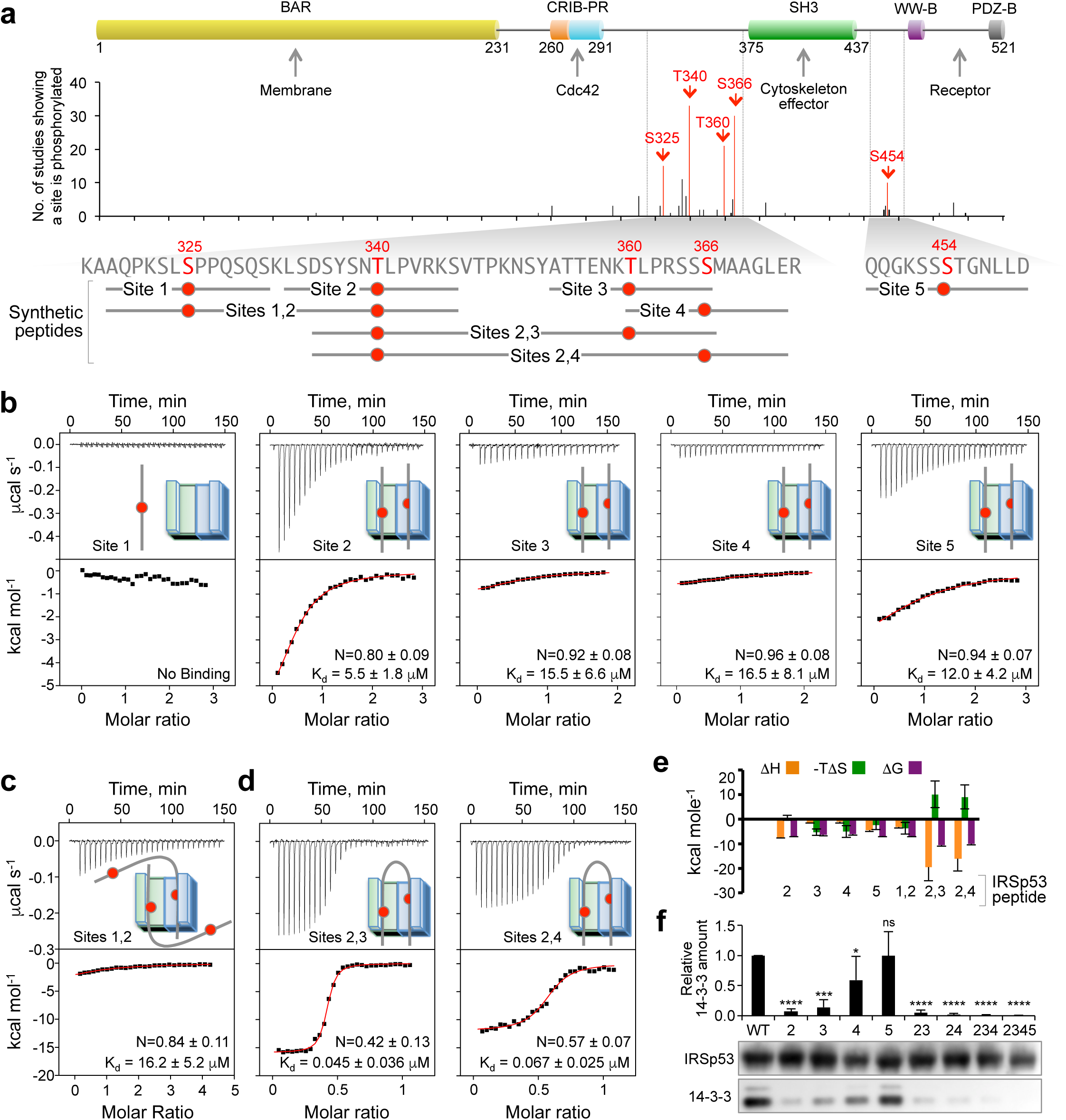
Characterization of the 14-3-3-binding sites of pIRSp53. **a** Domain organization of IRSp53, showing the location of phosphorylation sites and the definition of synthetic phospho-peptides used in this study. In the graph, the height of each phosphorylation site represents the number of times a particular residue has been found to be phosphorylated in phosphoproteomics studies curated by PhosphoSitePlus (www.phosphosite.org). Among these, five sites (highlighted red) were consistently found to be phosphorylated in four different samples analyzed here (purified as shown in **Fig. 1**). Note that the five sites identified here coincide with the highest frequency sites in PhosphoSitePlus. **b-d** ITC titrations of 100 μΜ singly-phosphorylated (Site 1 to 5) or 50 μΜ doubly-phosphorylated (Sites 1,2; 2,3 and 2,4) peptides into 10 μΜ 14-3-3θ (as indicated). ITC experiments were performed at 20°C. Listed for each experiment are the dissociation constant (K_d_) and binding stoichiometry (N) derived from fitting of the binding isotherm. Errors correspond to the s.d. of the fits. **e** Change in thermodynamic parameters derived from the fits of the ITC titrations shown in parts b-d. **f** Coimmunoprecipitation of 14-3-3 with WT IRSp53-FLAG and mutants carrying S/T->A substitutions of the indicated phosphorylation sites (named according to the number of the phosphorylation sites in part **a**. Error bars are ± s.d. from three independent experiments. The statistical significance of the measurements was determined using Student’s t-test, comparing each of the mutants to WT IRSp53 (ns, non-significant; *, p < 0.05; **, p < 0.01; ***, p < 0.001; ****, p < 0.0001).

Next, we asked which of these phosphorylation sites were implicated in 14-3-3 binding. Because the phospho-S/T-binding pockets of 14-3-3 are ~34 Å apart, it can in principle bind to either two identical sites on two subunits of a dimeric target or two distinct sites on a single polypeptide, in which case the sites are commonly found 19–40 amino acids apart within intrinsically disordered regions ^45^. Therefore, we used isothermal titration calorimetry (ITC) to test the phosphorylation sites identified here for their ability to bind *E. coli-expressed* 14-3-3θ individually and as pairs. Eight IRSp53 synthetic phospho-peptides were examined, including five 12-14-residue-long singly-phosphorylated peptides (named Site 1 to 5) and three 29–40-residue-long doubly-phosphorylated peptides (named Sites 1,2; 2,3 and 2,4) (**Fig. 2a** and **Supplementary Table 1**).

All the singly-phosphorylated peptides, except Site 1 (pS325), bound 14-3-3θ with ~1:1 stoichiometry (*i.e*. two peptides per 14-3-3θ dimer) and relatively low affinities (*K*_d_ raging from 5.5 to 16.5 μM) (**Fig. 2b,e**). The doubly-phosphorylated Sites 1,2 peptide (pS325/pT340) also bound 14-3-3θ with ~1:1 stoichiometry and micromolar affinity (**Fig. 2c,e**), suggesting that it interacts with 14-3-3θ only through Site 2 (pT340) thus ruling out Site 1 (pS325) as a potential 14-3-3θ-binding site. In contrast, the Sites 2,3 (pT340/pT360) and Sites 2,4 (pT340/pS366) peptides bound 14-3-3θ with ~0.5:1 stoichiometry *(i.e*. one peptide per 14-3-3θ dimer) and with more than two orders of magnitude higher affinities (*K*_d_ of 45 nM and 67 nM, respectively) than the corresponding singly-phosphorylated peptides (**Fig. 2d,e**). This result suggested that these doubly-phosphorylated peptides occupy the two phospho-S/T-binding pockets of 14-3-3θ. Of note, other targets of 14-3-3 also bind with nanomolar affinity and display similar increases in affinity for singly-phosphorylated *vs*. doubly-phosphorylated peptides ^45, 46, 47,48, 49, 50, 51^

A peptide encompassing pT360 and pS366 was not analyzed, because these two sites are too close to one another in the sequence to allow for simultaneous binding to the symmetric pockets of the 14-3-3 dimer. Site 5 (pS454), which alone bound 14-3-3θ with a *K*_d_ = 12 μM (**Fig. 2b**), could not be tested by ITC in combination with other sites, due to the inability to make synthetic peptides of the required length. However, a 4-alanine substitution of the ^452^SSST^455^ cluster had no effect on the relative amount of 14-3-3 that coimmunoprecipitates with IRSp53-FLAG from cells (**Fig. 2f**), allowing us to rule out this cluster as a 14-3-3-binding site. In contrast, single alanine substitutions of residues T340, T360 and S366 all significantly reduced 14-3-3 binding in cells, and any combination of these mutations completely abolished 14-3-3 binding (**Fig. 2f**). Importantly, phosphomimetic mutations of these sites cannot substitute for phosphorylation in the case of 14-3-3 binding; *E. coli-*expressed phosphomimetic mutants of full-length IRSp53 containing aspartic acid or glutamic acid at positions 340, 360, 366 and 454 failed to bind 14-3-3θ (**Supplementary Fig. 1**). From the ensemble of these results we conclude that 14-3-3 binds to three major sites in pIRSp53 (pT340, pT360 and pS366), and its binding affinity increases ~100-fold for two pairs of sites: pT340/pT360 and pT340/pT366. The existence of two pairs of 14-3-3-binding sites in pIRSp53 reconciles the apparent discrepancy between the findings of two previous studies ^35, 36^.

### Each subunit of the pIRSp53 dimer binds one 14-3-3 dimer

Both 14-3-3 and IRSp53 form constitutive dimers, as do several targets of 14-3-3, including p53, c-Raf, LRRK2, ASK1 and HSPB6. A recent crystal structure of HSPB6 in complex with 14-3-3σ ^52^ and a SAXS study of pASK1-CD in complex with 14-3-3ζ ^53^ show a single 14-3-3 dimer bound to a single phosphorylation site within one of the subunits of these dimeric targets. However, 14-3-3 binds to some targets through two or more phosphorylation sites, and whether 14-3-3 binds to different sites within a single subunit or to identical (or different) sites on two subunits of a dimeric target is a question that has not been formally addressed. We do this here for IRSp53, both biochemically and in cells.

In cells, a dual tag (FLAG, STREP) approach was developed for the purification of heterodimers of WT IRSp53 and a mutant in which residues T340, T360, S366 and ^452^SSST^455^ were simultaneously mutated to alanine (M2345). Importantly, a first attempt, using two separate plasmids to co-express WT IRSp53-FLAG and M2345-STREP in HEK293T cells (**Supplementary Fig. 2a**), failed to produce heterodimers (**Supplementary Fig. 2b**). This approach produced only homodimers (WT/WT or M2345/M2345), strongly suggesting that IRSp53 dimerization occurs during translation and is irreversible. In contrast, WT/M2345 heterodimers were successfully expressed from a single vector, coding a single transcription product. For this, we inserted an internal ribosomal entry site (IRES) between the WT and M2345 coding sequences, allowing for the simultaneous translation of two open reading frames (**Fig. 3a**). When both subunits were mutated, 14-3-3 did not coimmunoprecipitate with M2345/M2345, whereas approximately 50% of the amount that coimmunoprecipitates with WT/WT IRSp53 was observed when one of the dimer subunits was mutated (WT/M2345 or M2345/WT), independent of which affinity tag was attached to the mutant subunit or whether the mutant subunit preceded or followed the IRES (**Fig. 3b**). These results demonstrate that 14-3-3 binds to two different phosphorylation sites on a single subunit of the pIRSp53 dimer in cells.

**Fig. 3.**
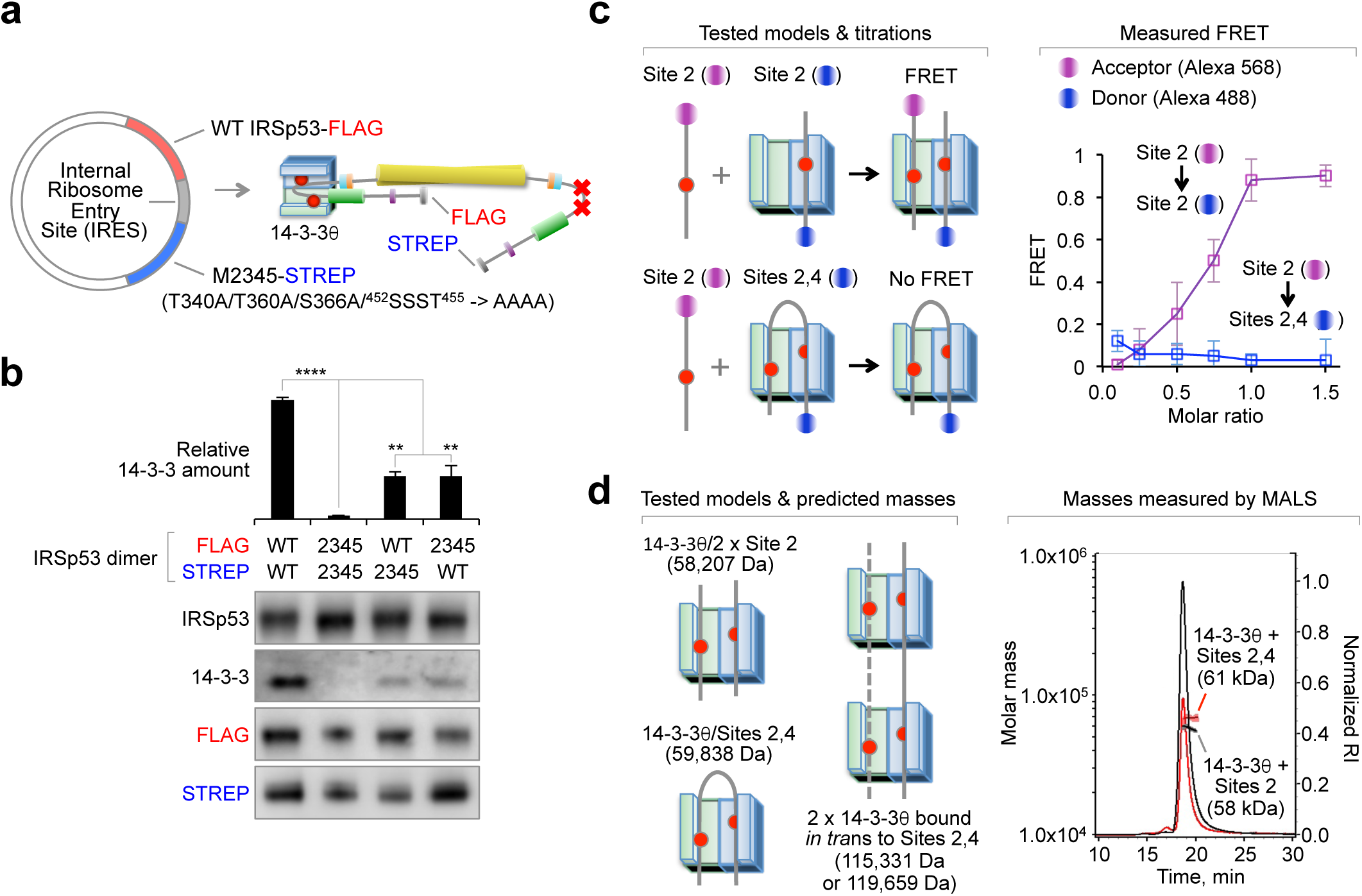
Each subunit of the pIRSp53 dimer binds one 14-3-3 dimer. **a** Illustration of the strategy used to obtain wild-type/mutant IRSp53 heterodimers using dual affinity purification tags (FLAG and STREP) and an internal ribosomal entry site (IRES) inserted between the wild-type and mutant coding sequences of a single expression plasmid. The mutant M2345 carries seven S/T->A substitutions (T340A/T360A/S366A/^452^SSST^455^ -> AAAA). **b** Coimmunoprecipitation of 14-3-3 with homo- and heterodimers of wild-type and mutant IRSp53. Two different heterodimers were tested, where the mutant subunit either preceded or followed the IRES, which also changed the purification tag. Error bars are ± s.d. from three independent experiments. The statistical significance of the measurements was determined using Student’s t-test, comparing each of the samples to wild-type IRSp53 (**, p < 0.01; ****, p < 0.0001). **c** FRET assay in which acceptor (Alexa 568)-labeled Site 2 peptide was titrated into 14-3-3θ half-saturated (1:0.5 molar ratio) with donor (Alexa 488)-labeled Site 2 or Sites 2,4 peptides. FRET is only observed when the acceptor-labeled peptide binds to one of the pockets of the 14-3-3θ dimer while the other pocket is occupied by donor-labeled peptide. Error bars are ± s.d. from three independent titrations. **d** Determination of the masses of complexes of 14-3-3θ with Site 2 or Sites 2,4 peptides using SEC-MALS.

To further validate the interaction of 14-3-3 with pairs of sites within a single subunit of the pIRSp53 dimer, a FRET competition assay was developed to monitor the binding of acceptor-labeled Site 2 peptide to half-saturated (1:0.5 stoichiometry) complexes of 14-3-3θ with either donor-labeled Site 2 or Sites 2,4 peptides (**Fig. 3c**). The titration into the half-saturated 14-3-3θ/Site 2 complex produced a large FRET increase, indicating that the acceptor-labeled Site 2 peptide bound to the vacant pocket of the 14-3-3θ dimer, within FRET distance from the donor-labeled Site 2 peptide (pre-bound to the other pocket of 14-3-3θ). In contrast, the titration of acceptor-labeled Site 2 peptide into the 14-3-3θ/Sites 2,4 complex did not produce a FRET change, suggesting that both pockets of 14-3-3θ were occupied by the doubly-phosphorylated peptide, and that the singly-phosphorylated peptide cannot outcompete the doubly-phosphorylated peptide from either pocket.

A doubly-phosphorylated peptide could in principle bind to two 14-3-3θ dimers in *trans*, a possibility tested here using multi-angle light scattering to measure the masses of saturated complexes of 14-3-3θ with singly- and doubly-phosphorylated peptides (**Fig. 3d**). The experimentally determined mass of the 14-3-3θ/Site 2 complex was 58,000 Da, consistent with a 1:1 complex (theoretical mass: 58,207 Da), whereas that of 14-3-3θ/Sites 2,4 was 61,000 Da, corresponding to a 1:0.5 complex (theoretical mass: 59,838 Da). These masses rule out the formation of 14-3-3θ complexes in *trans* with the doubly-phosphorylated Sites 2,4 peptide, with theoretical masses of 115,331 Da (one peptide bound *in trans* to two 14-3-3θ dimers) or 119,659 Da (two peptides bound *in trans* to two 14-3-3θ dimers). Together, these results suggest that 14-3-3 binds to two pairs of sites (pT340/pT360 or pT340/pS366) within a single subunit of the IRSp53 dimer.

### Structural characterization of the interaction of 14-3-3 with pIRSp53

Three factors make the structural characterization of complexes of 14-3-3 with doubly-phosphorylated peptides challenging: a) the low-complexity regions between phosphorylation sites are usually disordered in the structures of 14-3-3 complexes, b) 14-3-3 binds to phospho-S/T sites that are related in sequence and thus difficult to tell apart in electron density maps, and c) doubly-phosphorylated peptides can in principle bind with two opposite N- to C-terminal orientations, such that each of the 14-3-3 pockets could in principle contain a mixture of the two phosphorylation sites present in the peptide. Therefore, to study the binding of 14-3-3θ to the pT340/pT360 and pT340/pS366 pairs, we determined five crystal structures, allowing for direct comparisons of the structures of singly-*vs*. doubly-phosphorylated complexes (**Fig. 4** and **Supplementary Fig. 3**). In the structures, ranging in resolution from 2.0 to 2.9 Å (**Table 1**), the electron density maps revealed the phosphorylated residue and 3 to 5 residues N- and C-terminal to it, whereas the ends of the peptides and the flexible linkers between phosphorylation sites in the doubly-phosphorylated peptides were unresolved. In all the structures, the phosphate group on the phosphorylated side chain engages a cluster of basic residues in 14-3-3θ, consisting of K49, R56, and R127, and additionally forms a hydrogen bond with the hydroxyl group of Y128. For the singly-phosphorylated peptides, the electron density is symmetric for the two pockets of 14-3-3θ, even when two-fold symmetry was not imposed during refinement. Importantly, the structures of the singly-phosphorylated peptides display distinctive features that are recognizable in those of the doubly-phosphorylated peptides, for which the electron density is asymmetric in the two pockets of 14-3-3θ (**Fig. 4**). Specifically, the Site 2 and Site 3 peptides contain a proline residue at the second position after the phosphorylation site, producing a characteristic 90° bend of the polypeptide chain, which as a result exits the 14-3-3 binding groove, thus leaving a significant portion of the groove unoccupied (**Supplementary Fig. 3a,b**). In contrast, the Site 4 peptide has an alanine residue at the second position after the phosphorylation site (pS366), and follows a relatively straight path along the groove of 14-3-3θ (**Supplementary Fig. 3c**). Although the sequences surrounding pT340 and pT360 are closely related to each other, they are distinguishable by the presence of two large side chains in the Site 1 peptide, Y337 and R344, occupied by smaller side chains in the Site 2 peptide, E357 and S365, respectively (**Supplementary Fig. 3a,b**). These distinctive features of the singly-phosphorylated peptides are clearly discernable in the structures of the doubly-phosphorylated peptides (**Fig. 4a,b**), in which each pocket of 14-3-3θ binds a different site in an asymmetric manner. We note, however, that the structure of the pT340/pT360 pair contains four complexes in the P1 unit cell, and only one of these complexes has a unique N- to C-terminal orientation of the doubly-phosphorylated peptide, whereas the other three complexes appear to have a mixture of orientations. Together, the biochemical (**Figs. 2** and **3**) and structural (**Fig. 4**) data allow us to conclude that 14-3-3 binds with high affinity to two pairs of sites in pIRSp53, pT340/pT360 and pT340/pS366.

**Fig. 4.**
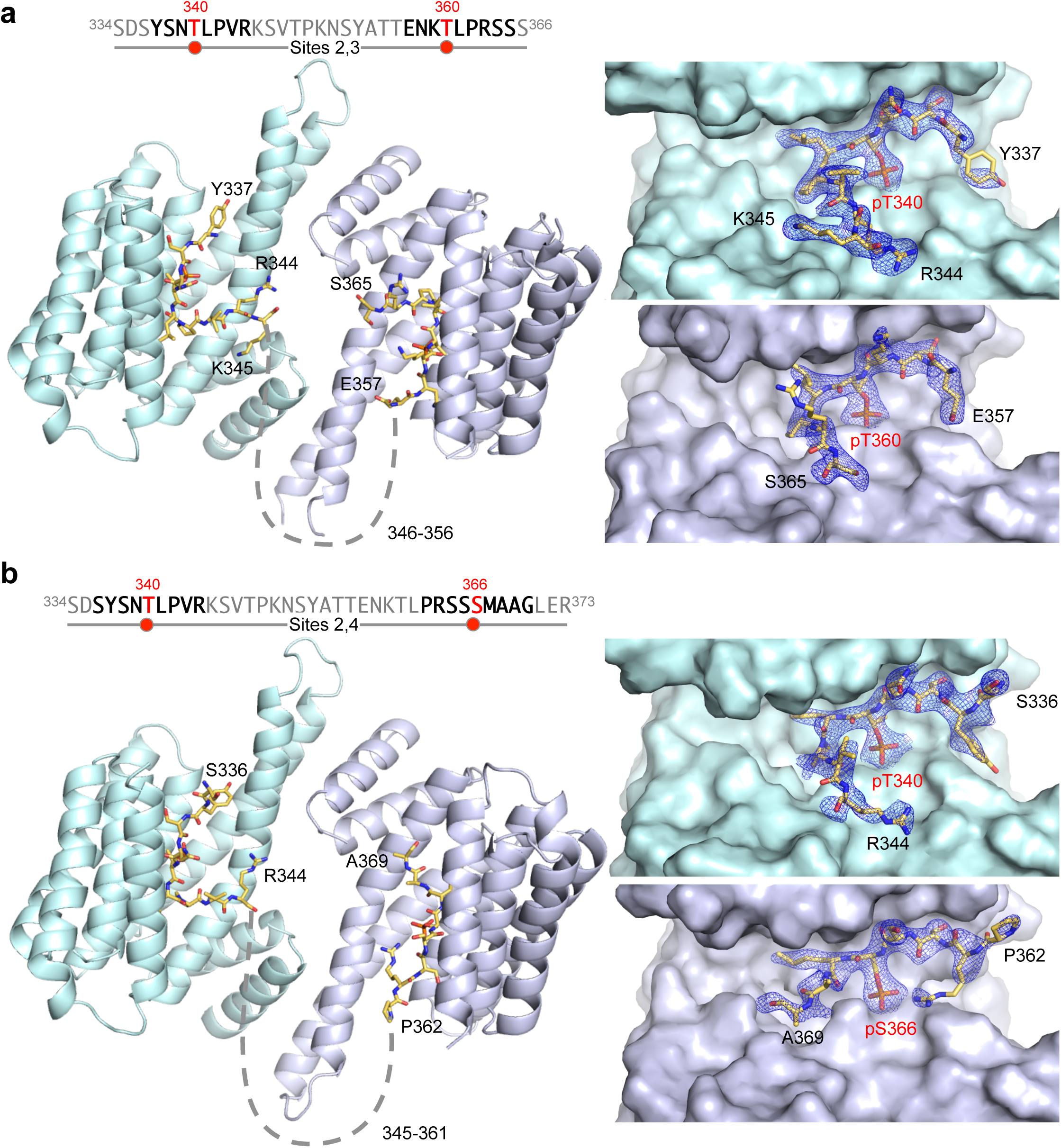
Structures of complexes of 14-3-3θ with doubly-phosphorylated IRSp53 peptides. **a,b** Complexes of 14-3-3θ with the Sites 2,3 and Sites 2,4 peptides, showing the overall structure (left) and close-ups of the binding pockets of 14-3-3θ and corresponding 2F_o_-F_c_ electron density maps contoured at 1 σ (right). In the sequence diagrams shown, the portions of the peptides observed in the structures are highlighted bold and the phosphorylation sites are highlighted red. Dashed lines indicate the path between the observed portions of the bound peptides. Note that the sequences around the two phosphorylation sites of each peptide are different enough to be clearly distinguishable in the electron density maps in the separate pockets of 14-3-3θ, as also confirmed by cross-comparisons with the structures of the corresponding singly-phosphorylated peptides (**Supplementary Fig. 2**).

**Table 1.** Crystallographic data and refinement statistics

| Table 1. Crystallographic data and refinement statistics |  |  |  |  |  |
| --- | --- | --- | --- | --- | --- |
| Structure | Site 2 (pT340) | Site 3 (pT360) | Site 4 (pSS366) | Sites 2,3 (pT340/pT360) | Sites 3,4 (pT340/pSS366) |
| **Data collection** |  |  |  |  |  |
| Beamline | NSLS X6A | MacCHESS A1 | X8 Prospector | X8 Prospector | X8 Prospector |
| Wavelength (Å) | 1.000 | 0.977 | 1.540 | 1.540 | 1.540 |
| Space group | P 1 | P 1 | P 1 | P 1 | P 21 21 21 |
| Cell *a*, *b*, *c* (Å) | 60.0, 69.1, 84.6 | 59.4, 68.6, 84.8 | 59.9, 69.3, 84.7 | 61.1, 69.6, 150.5 | 69.9, 75.0, 100.2 |
| Cell $\alpha$, $\beta$, $\gamma$ (°) | 105.3, 95.7, 115.0 | 72.9, 85.6, 64.7 | 106.1, 95.6, 114.9 | 99.2, 93.0, 116.0 | 90.0, 90.0, 90.0 |
| Resolution (Å) | 31.3–2.0 (2.06–2.0)<sup>a</sup> | 21.9–2.3 (2.38–2.3) | 29.4–2.35 (2.4–2.35) | 43.0–2.8 (2.9–2.8) | 60.0–2.9 (3.0–2.9) |
| *R*<sub>merge</sub> (%) | 4.2 (37.9) | 8.7 (57.9) | 5.8 (20.5) | 14.2 (29.3) | 27.8 (34.4) |
| *I* / σ*I* | 21.8 (3.5) | 12.5 (3.6) | 20.0 (4.9) | 20.5 (2.5) | 17.0 (2.1) |
| No. unique reflections | 76071 (6527) | 49462 (4819) | 45896 (4330) | 51911 (5197) | 12087 (1064) |
| Multiplicity | 3.4 (2.7) | 3.6 (3.2) | 5.5 (2.7) | 4.8 (2.1) | 7.2 (1.9) |
| Completeness (%) | 95.7 (92.0) | 96.6 (94.3) | 95.6 (90.4) | 96.7 (95.7) | 98.8 (91.4) |
| CC<sub>1/2</sub> | 0.999 (0.888) | 0.996 (0.87) | 0.998 (0.889) | 0.975 (0.717) | 0.887 (0.629) |
| Wilson B-factor (Å<sup>2</sup>) | 32.75 | 35.95 | 23.21 | 30.91 | 25.61 |
| **Refinement** |  |  |  |  |  |
| Resolution (Å) | 31.3–2.0 | 21.9–2.3 | 29.4–2.35 | 43.0–2.8 | 60.0–2.9 |
| No. reflections | 76054 | 49428 | 45857 | 51490 | 12042 |
| *R*<sub>work</sub> (%) / *R*<sub>free</sub> (%) | 19.5 / 22.9 | 19.2 / 22.2 | 21.2 / 25.1 | 23.5 / 28.6 | 21.5 / 25.8 |
| No. atoms |  |  |  |  |  |
| Protein | 7868 | 7763 | 7692 | 15360 | 3868 |
| Ligands | 92 | 111 | 70 | 212 | 50 |
| Solvent (H<sub>2</sub>O) | 517 | 110 | 406 | 93 | 34 |
| RMS deviations |  |  |  |  |  |
| Bond lengths (Å) | 0.003 | 0.002 | 0.002 | 0.005 | 0.004 |
| Bond angles (°) | 0.55 | 0.46 | 0.45 | 0.62 | 0.88 |
| B-factors (Å<sup>2</sup>) |  |  |  |  |  |
| Protein | 42.2 | 54.0 | 31.7 | 38.3 | 22.5 |
| Ligand | 52.4 | 60.9 | 40.6 | 46.5 | 34.5 |
| Solvent | 42.6 | 43.0 | 29.9 | 18.5 | 14.6 |
| Ramachandran (%) |  |  |  |  |  |
| Favored | 99.1 | 98.6 | 98.9 | 97.3 | 98.3 |
| Outliers | 0.0 | 0.0 | 0.0 | 0.0 | 0.0 |
| PDB code | 6BCR | 6BCY | 6BD1 | 6BQT | 6BD2 |

<sup>a</sup> Values in parentheses are for highest-resolution shell

### A FRET-sensor assay shows inhibition of pIRSp53 by 14-3-3

We then asked whether the binding of 14-3-3 to pIRSp53 regulates its activity by stabilizing the closed, autoinhibited conformation in which an intramolecular interaction between the CRIB-PR and SH3 domains blocks access to these two domains ^10^. To test this idea, a FRET-sensor assay was developed to monitor conformational changes in pIRSp53 as a function of its interaction with binding partners. This approach builds upon our previous report of a FRET-sensor using *E. coli*-expressed IRSp53 ^10^, but we now use the purification protocol described here (**Fig. 1**) to obtain a FRET sensor natively phosphorylated in mammalian cells (pIRSp53_FS_). The protein used in the FRET-sensor assay was labeled with the donor/acceptor pair fluorescein/rhodamine at two positions: endogenous C230 within the BAR domain (the only label-accessible Cys residue in IRSp53) and C519 introduced by mutagenesis near the C-terminus (S519 in WT IRSp53). Conformational changes in pIRSp53_FS_ were monitored as the change in energy transfer resulting from the binding of several inputs: 14-3-3θ, Cdc42 and the cytoskeletal effector Eps8 (**Fig. 5a**). All the proteins used in these experiments were purified to homogeneity (**Supplementary Fig. 4**).

**Fig. 5.**
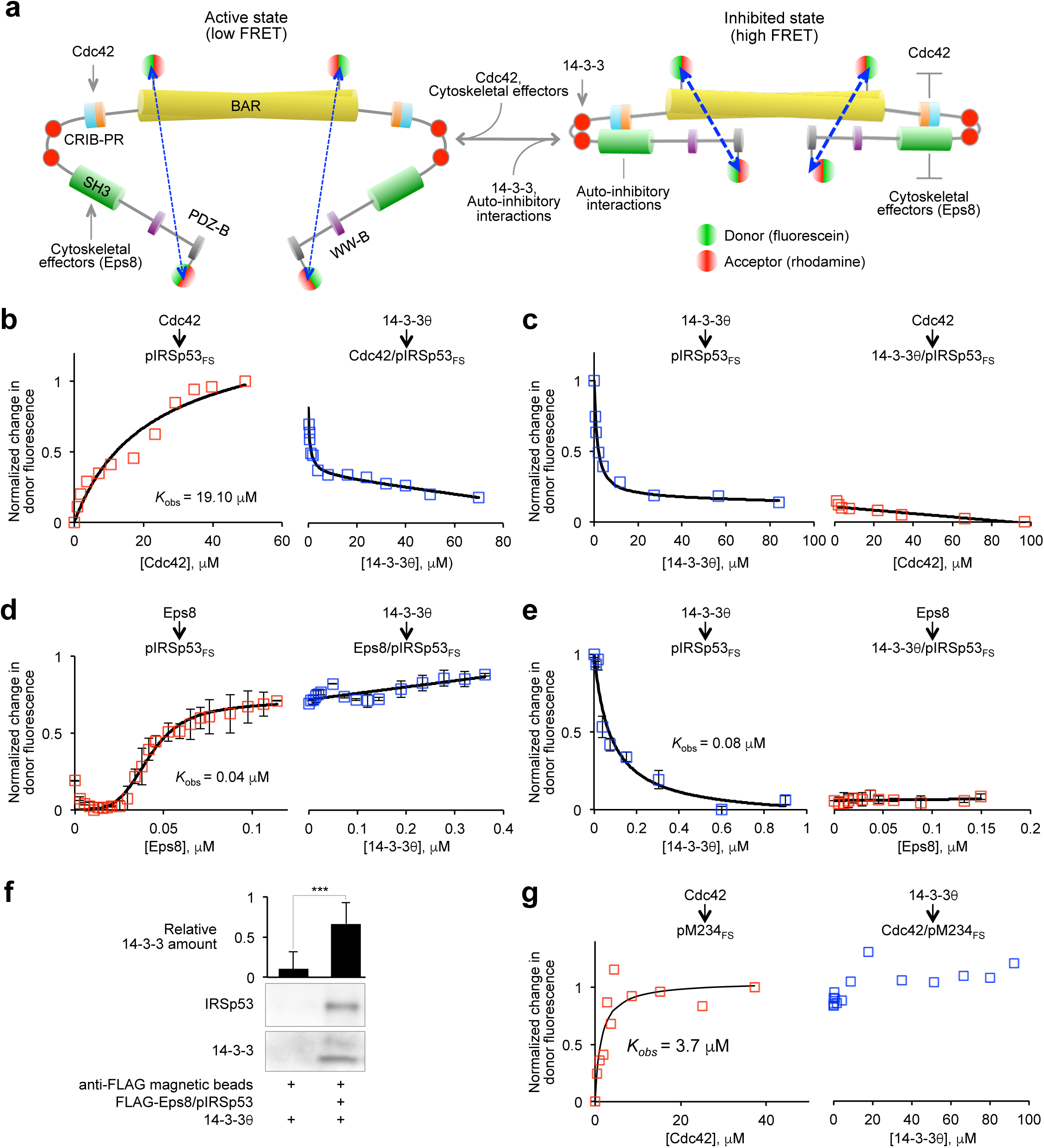
A FRET-sensor assay shows inhibition of pIRSp53 by 14-3-3. **a** FRET sensor assay designed to report on conformational changes occurring in pIRSp53 as a function of interactions with binding partners. Phosphorylated IRSp53 (mutant S519C) was expressed and purified as shown in Fig. 1 and labeled at C230 (within the BAR domain) and C519 (near the C-terminus) with the donor/acceptor pair fluorescein/rhodamine (pIRSp53_FS_). The same batch of probes and donor/acceptor ratio were used in the labeling reactions. Conformational changes in pIRSp53_FS_ can be detected based on the amount of FRET between fluorescence probes at these two distant positions. **b**,**c** FRET titration of 2 μΜ GNPPNP-Cdc42 (G12V mutant) into 0.2 μΜ pIRSp53_FS_, followed by the titration of 0.2 μΜ 14-3-3θ and reverse-order titration. **d**,**e** FRET titration of 0.2 μΜ Eps8 into 0.05 μΜ pIRSp53_FS_, followed by the titration of 4 μΜ 14-3-3θ and reverse-order titration. Pairs of titrations were normalized to the total change in FRET. Each titration was fit to a non-cooperative binding function (black lines), except that of Eps8 into pIRSp53_FS_, which was fit to a cooperative binding function with non-specific binding (see Methods). Error bars are ± s.d. from three independent experiments. **f** Relative abundance of 14-3-3 pulling down with anti-Flag magnetic beads in the presence or the absence of the FLAG-Eps8/pIRSp53 complex. Error bars are ± s.d. from three independent experiments. The statistical significance of the measurements was determined using Student’s t-test (***, p < 0.001). **g** FRET titration of 2 μΜ GNPPNP-Cdc42 into 0.2 μΜ mutant Μ234|γ_5_ of IRSp53 (purified as shown in Fig. 1), followed by the titration of 0.2 μΜ 14-3-3.

The titration of GMPPNP-Cdc42 (constitutively active mutant G12V) into pIRSp53_FS_ induced a transition toward an open, active conformation, characterized by an increase in the donor fluorescence *(i.e*. a low FRET state) (**Fig. 5b**). The subsequent titration of 14-3-3θ into this complex reduced the donor fluorescence (high FRET state), consistent with a reversal of the conformational change induced by Cdc42 and a transition of pIRSp53 toward a closed, inactive conformation. When the order of the titrations was inverted, 14-3-3θ triggered a transition of pIRSp53_FS_ toward a closed conformation, which remained mostly unchanged with the subsequent titration of Cdc42 (**Fig. 5c**). The reason for the small increase in energy transfer upon titration with Cdc42 is unclear, but could indicate non-activatory binding of Cdc42 to the IRSp53-14-3-3 complex. These results indicate that Cdc42 cannot competitively activate pIRSp53 once it is bound to 14-3-3.

Since, the binding of cytoskeletal effectors such as Eps8 to the SH3 domain also activates IRSp53 ^10^, we performed back-to-back titrations of 14-3-3θ and Eps8 into pIRSp53_FS_, using Eps8 purified from mammalian cells (**Supplementary Fig. 4**). The binding of Eps8 to pIRSp53_FS_ increased the donor fluorescence in a cooperative manner (**Fig. 5d**), consistent with a transition of pIRSp53 toward an open, active state. As we have shown before ^10^, the Eps8/pIRSp53 complex has 2:2 stoichiometry. However, the subsequent titration of 14-3-3θ into this complex did not revert the conformational change produced by Eps8. On the contrary, the donor fluorescence increased linearly, albeit only by ~15% with a 4-fold molar excess of 14-3-3θ to Eps8 (**Fig. 5d**). This effect is possibly explained by the inability of 14-3-3θ to displace Eps8 from its complex with pIRSp53, as also suggested by the observation that FLAG-tagged Eps8 pulls down both pIRSp53 and 14-3-3θ (**Fig. 5f**). We thus conclude that the binding of 14-3-3θ to Eps8/pIRSp53 produces a low-FRET ternary complex, whose conformation is distinct from those of the active Eps8/pIRSp53 and the inhibited 14-3-3θ/pIRSp53 complexes, and which we cannot reliable classify as either open or closed. The ability of 14-3-3θ to competitively revert the conformational change induced by Cdc42 but not Eps8 is consistent with the relative binding affinities of these three proteins for pIRSp53; from fitting of the titrations the observed *K_d_* (*K_obs_*) of the interactions of pIRSp53 with Cdc42, Eps8 and 14-3-3θ were estimated at 19.1 μM, 0.042 μM and 0.079 μM, respectively. We then inverted the order of the titrations; 14-3-3θ induced the expected transition of pIRSp53 toward a closed, inactive state, and the subsequent titration of Eps8 into this complex did not change the donor fluorescence (**Fig. 5e**), suggesting that even a high-affinity cytoskeletal effector such as Eps8 cannot competitively activate pIRSp53 once it is bound to 14-3-3.

To test whether the effect of 14-3-3θ on the conformation of pIRSp53 depends on binding to the two pairs of phosphorylation sites identified above, we expressed in serum-starved HEK293T cells a modified version of the FRET sensor (pM234_FS_) in which these sites were simultaneously mutated to alanine (T340A/T360A/S366A). As shown above (**Fig. 2f**), this mutant fails to coimmunoprecipitate with 14-3-3 from cells. The titration of Cdc42 into pM234_FS_ induced the anticipated transition to an open conformation, but contrary to pIRSp53_FS_ the subsequent titration of 14-3-3θ did not revert this conformational change (compare **Figs. 5b** and **5g**). Together, these results suggest that 14-3-3 “locks” pIRSp53 in a closed, autoinhibited conformation, blocking the interactions of Cdc42 and cytoskeletal effectors with the CRIB-PR and SH3 domains.

### Binding of 14-3-3 reduces pIRSp53’s association with membranes

We then asked whether the binding of 14-3-3 inhibits the interaction of pIRSp53 with membranes in cells. To this end, we separated the cytosolic and membrane fractions of fed and serum-starved HEK293T cells expressing either WT IRSp53-FLAG or the mutant M234-FLAG (unable to bind 14-3-3), and determined the relative abundance of the two IRSp53 constructs in the two cellular fractions. Independent of starvation, IRSp53-FLAG was more abundant in the cytosolic than the membrane fraction, whereas the opposite was observed with M234-FLAG, which was more abundant in the membrane fraction (**Fig. 6a**). Furthermore, serum starvation, a condition that increases the amount of 14-3-3 that coimmunoprecipitates with FLAG-IRSp53 (**Fig. 1a**), reduced the amount of WT IRSp53-FLAG bound to membranes by ~10% compared to fed cells, but had no significant effect on the distribution of M234-FLAG between the cytosolic and membrane fractions. We thus conclude that 14-3-3 not only inhibits the binding of pIRSp53 to Cdc42 and cytoskeletal effectors, but also interferes with its interaction with cellular membranes.

**Fig. 6.**
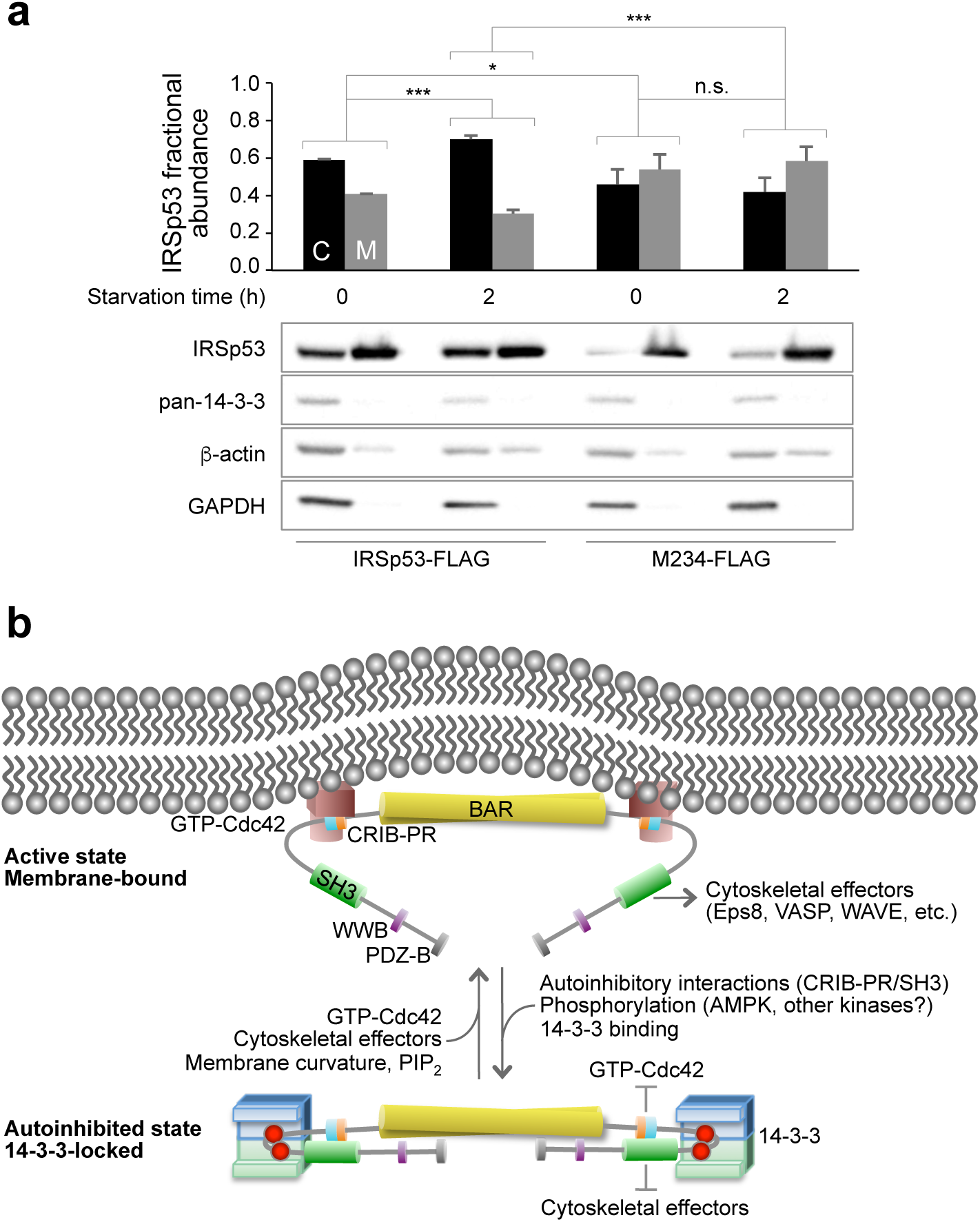
Mechanism of IRSp53 regulation by 14-3-3. **a** 14-3-3 inhibits membrane binding of pIRSp53. Relative abundance of WT IRSp53-FLAG and mutant M234-FLAG in the cytosolic (C) and membrane (Μ) fractions of fed and serum-starved HEK293T cells. The statistical significance of the measurements was determined using Student’s t-test, based on data from three independent transfections per condition (n.s., non-significant; *, p < 0.05; **, p < 0.01; ***, p < 0.001). Error bars are ± s.d. A representative Western blot analysis is shown, using GAPDH and β-actin as loading controls. **b** Model for IRSp53 activation and phosphorylation-dependent inhibition by 14-3-3. IRSp53 is activated by the binding of Cdc42 to the CRIB-PR and/or cytoskeletal effector proteins to the SH3 domain. Activation also exposes the I-BAR domain for binding to the plasma membrane. Phosphorylation of IRSp53 and the subsequent binding of 14-3-3 to phosphorylation sites located between the CRIB-PR and SH3 domains stabilizes the autoinhibited conformation and blocks the binding of Cdc42 and downstream cytoskeletal effectors.

## Discussion

Membrane remodeling during processes such as tissue development and cancer cell migration depends on the interplay between membrane-associated proteins and the actin cytoskeleton. The I-BAR domain protein IRSp53 is a key player in these processes, acting as a hub that integrates signaling cues from Rho-GTPases and phosphorylation pathways to initiate cytoskeleton and membrane remodeling events ^2, 7^. 14-3-3 proteins, on the other hand, are a conserved family of phospho-S/T adaptors, known primarily for their roles in the control of cell cycle progression and apoptosis ^54^ and the regulation of cell metabolism ^55^. Yet, 14-3-3 has been also implicated in interactions with a growing number of actin cytoskeletal proteins ^35, 36, 39, 56, 57, 58, 59^, and it is known to impact both membrane morphology and actin cytoskeleton assembly ^56, 57^, although the mechanism is not well understood. This study provides a potential mechanism by linking 14-3-3 to the regulation of pIRSp53.

Using a combination of structural, biochemical and biophysical approaches, this study supports a mechanism whereby phosphorylation-dependent inhibition of pIRSp53 by 14-3-3 counters activation by Cdc42 and downstream cytoskeletal effectors and binding to cellular membranes (**Fig. 6b**). The major findings in support of this conclusion are: ***a***) a combination of phosphoproteomics, pull-down assays, ITC, and competitive binding analysis by FRET and light scattering of doubly-phosphorylated *vs*. singly-phosphorylated IRSp53 peptides demonstrates that 14-3-3 binds to two pairs of sites in the linker between the CRIB-PR and SH3 domains of pIRSp53 (pT340/pT360 or pT340/pS366), ***b***) analysis of an IRSp53 WT/M2345 heterodimer in which one subunit of the dimer is non-phosphorylatable shows that 14-3-3 binds to pairs of sites within a single subunit of the IRSp53 dimer, ***c***) the interaction of 14-3-3θ with the phosphorylation sites of pIRSp53 was visualized in five crystal structures of singly- and doubly-phosphorylated IRSp53 peptides bound to 14-3-3θ, ***d***) a FRET sensor assay based on natively phosphorylated IRSp53 purified from mammalian cells shows that 14-3-3 locks pIRSp53 in a closed, autoinhibited conformation, whereby access of Cdc42 to the CRIB-PR and cytoskeletal effectors to the SH3 domain are both inhibited, ***e***) membrane fractionation shows greater association of mutant M234 with cellular membranes than WT IRSp53, and partial depletion of WT IRSp53 but not M234 from membranes upon serum starvation.

Some of the experimental approaches developed here may find general applicability in the study of other 14-3-3 interactions, and in particular the FRET sensor assay. Indeed, the binding of 14-3-3 often results in a conformational change that either inhibits or unleashes a new function in the target protein by masking or exposing functional domains ^60, 61^. In such cases, it is essential to understand how the binding of 14-3-3 affects the structure and interactions of its targets, for which FRET sensors provide a simple and broadly viable tool. FRET sensors may also be used to identify novel modulators of 14-3-3 interactions, which may have potential therapeutic applications ^62^.

There are several isoforms of IRSp53, differing primarily at the C-terminus. In this study we focused on isoform-4, which is exclusively expressed in brain, and has a PDZ-binding site at the C-terminus^63^. The 14-3-3-binding sites identified here (pT340, pT360 and pS366) are conserved among all the isoforms, and among different species, suggesting that the mechanism of 14-3-3 regulation described here is conserved. In the future, it would be interesting to explore the ability of 14-3-3 to regulate other members of the I-BAR family, which are generally related to IRSp53 and often linked to cytoskeleton remodeling. Algorithms that predict potential 14-3-3-binding sites could help guide such studies. In this regard, we note that among several prediction servers tested, 14-3-3-Pred, which combines three different classifiers to predict 14-3-3-binding sites ^64^, showed the best performance. It identified five potential 14-3-3-binding sites within the region between the CRIB-PR and SH3 domains of IRSp53 (residues 292-374): S303, T340, T348, T360, and S366. This prediction includes the three major sites identified here: T340 and T360 (predicted by two classifiers) and S366 (predicted by three classifiers).

Because the 14-3-3 dimer contains two phospho-S/T-binding pockets, it could in principle bind to two ligands simultaneously, and function as a protein-protein adaptor. Yet, such mode of binding is rare, as the majority of 14-3-3 ligands contain two or more 14-3-3-binding sites that simultaneously engage the two pockets of the 14-3-3 dimer^45, 46, 61^. Therefore, our finding here that 14-3-3 inhibits pIRSp53 by binding to two different pairs of phosphorylation sites is far from unique, and instead suggests the existence of alternate routes of IRSp53 regulation, likely involving different kinases. The two pairs of 14-3-3-binding sites in pIRSp53 have one site in common, pT340, suggesting that pT360 and pS366 act as secondary sites in these pairs. This mode of action is consistent with the “gatekeeper” model, according to which phosphorylation of a first, high affinity site leads to the initial recruitment of 14-3-3 to a multi-site ligand, followed by phosphorylation and binding of the second pocket of 14-3-3 to a weaker secondary site, which in turn triggers a conformational change in the target protein ^60^. Coincidentally, 14-3-3 binds with 3-fold higher affinity to pT340 than to pT360 or pS366, whereas its affinity for the two pairs of sites is ~100-fold higher.

The identity of the kinases implicated in IRSp53 phosphorylation, and specifically at 14-3-3-binding sites, is mostly unknown. The sole exception is AMPK and AMPK-related kinases, which studies using chemical genetics screening have conclusively implicated in direct phosphorylation of IRSp53 at S366 ^35, 41, 42^. Furthermore, one of these studies linked the phosphorylation of this site to the binding of 14-3-3, with the concomitant loss of cell polarity and cell spreading in Madin–Darby canine kidney (MDCK) cells ^35^. On the other hand, the kinases implicated in the phosphorylation of T340 and T360 have not been identified, although GSK3β has been suggested to indirectly influence the phosphorylation of these two sites, based on the observation that LiCl, a potent and specific inhibitor of GSK3β, significantly reduces binding of 14-3-3 to IRSp53^36^. Two other kinases, LRRK2 ^65^ and PKD ^66^, have been indirectly implicated in changes in T340 phosphorylation, although none has been shown to directly phosphorylate this site. Interestingly, like AMPK-related kinases, PKD works hand-in-hand with 14-3-3 in other cytoskeleton regulatory pathways. Thus, PKD-mediated phosphorylation of cortactin stimulates 14-3-3 binding, which interferes with cortactin’s role as a stabilizer of Arp2/3 complex-mediated actin branches ^67^. PDK also phosphorylates the cofilin phosphatase slingshot-1L (SSH1L) to generate a 14-3-3-binding motif in SSH1L, which enhances the phosphorylation of cofilin and inhibits directed cell migration ^68^. In the future it would be important to conclusively identify the kinases involved in IRSp53 regulation at 14-3-3-binding sites, and understand how they synergize with AMPK-related kinases to influence IRSp53’s numerous activities and diverse interactions with downstream cytoskeletal effectors.

## Methods

### Protein cloning, expression and purification

For Co-IPs and protein purification, full-length IRSp53 (UniProt: Q9UQB8-4) was PCR-extended to include C-terminal FLAG- or STREP-affinity tags and cloned into vector pEGFP-C1 using the Nhel and Sall restriction sites. IRSp53 mutations T340A, T360A, S366A and S519C were introduced using the QuickChange II XL mutagenesis kit (Agilent Technologies, Santa Clara, CA). The ^452^SSST^455^ to ^452^AAAA^455^ mutation was generated by overlap PCR (primers listed in **Supplementary Table 2**). For the expression of wild-type/mutant IRSp53 heterodimers, the first and second genes were cloned respectively into the NheI/EcoRI and BamHI/NotI restriction sites of a modified vector pEGFP-C1 (containing a unique NotI restriction site in the multi-cloning site), carrying an internal ribosomal entry site (IRES) that was amplified from the plasmid pEF1a-IRES-Neo (Addgene, Cambridge, MA) and inserted into the EcoRI/BamHI restriction sites. This strategy allowed for the translation of two open reading frames from a single messenger RNA. For the expression of 2xFLAG-Eps8, full-length human Eps8 (UniProt: Q12929) was cloned into vector pEGFP-C1 and an oligonucleotide encoding for the sequence MDYKDHDGDYKDDDDKG was inserted between the NheI and EcoRI sites.

#### Mammalian cell protein expression and purification

For the expression of the natively phosphorylated IRSp53-FLAG used in the FRET sensor assay, HEK293T cells (ATCC, Manassas, VA) were grown in DMEM GlutaMAX medium supplemented with 5% FBS and antibiotic-antimycotic (Thermo Fisher Scientific, Waltham, MA) to ~40% confluence in 15-cm plates and transiently transfected with 10 μg plasmid DNA in 30 of a 1 mg mL^−1^ polyethylenimine (PEI) solution (Sigma-Aldrich Corporation, St. Louis, MO). After 48 h expression, IRSp53 phosphorylation was increased by serum-depriving cells for 4 h. Cells were then washed and incubated in PBS with the addition of 0.1 μM Calyculin A (Cell Signaling Technology, Danvers, MA) and 1x Halt Phosphatase Inhibitor Cocktail (Thermo Fisher Scientific) for 15 min at 37°C. Cells were harvested, pelleted and resuspended in lysis buffer comprised of PBS supplemented with 1 mM EDTA, 5% Glycerol, 1% Triton X100, 1 mM PMSF, 0.1 μM Calyculin A, 1x cOmplete protease inhibitor cocktail (Sigma-Aldrich Corporation) and 1x Halt Phosphatase Inhibitor Cocktail. Cells were incubated in lysis buffer on ice for 30 min and freeze-thawed three-times. Insoluble cellular components and nuclei were removed by centrifugation at 12,000g for 10 min. Clarified lysates were incubated with 1 mL anti-FLAG antibody resin (Thermo Fisher Scientific) for 2 h at 4°C and washed with 20 column volumes of lysis buffer (without Triton X100). Endogenous 14-3-3 was removed by washing the resin with 5 column volumes of wash buffer (PBS, 1 mM EDTA, 5% glycerol, 1x cOmplete protease inhibitor cocktail, and 1x Phosphatase Inhibitor Cocktail) supplemented with 200 mM Sites 2,3 peptide. The excess peptide was removed with 10 column volumes of wash buffer, and IRSp53 eluted with wash buffer supplemented with 300 mM NaCl and 0.5 mg mL^−1^ 3xFLAG peptide (Thermo Fisher Scientific). The fraction of phosphorylated IRSp53 able to bind 14-3-3 was then isolated by loading onto a 14-3-3-affinity column, consisting of MBP-tagged 14-3-3θ (see expression below) bound to an amylose resin. The bound IRSp53 was eluted using 200 μM Sites 2,3 peptide, and dialyzed into 25 mM Tris-HCl pH 8.0, 150 mM NaCl, 5% glycerol and 1 mM DTT. The protein was then incubate for 30 min at 4°C with 200 μL of Macro-Prep High S cation exchange resin (Bio-Rad Laboratories, Hercules, CA). The resin was washed with 10 column volumes of dialysis buffer, followed by 5 column volumes of dialysis buffer supplemented with 200 mM NaCl. IRSp53 was eluted with 500 μL of 25 mM HEPES pH 7.5, 500 mM NaCl, 1 mM MgCl_2_, 5% glycerol.

The construct 2xFLAG-Eps8 was transfected and expressed in HEK293T cells in a similar manner, except cells were not serum-starved or treated with phosphatase inhibitors. Cell lysis and FLAG-affinity purification were also performed as described above, except phosphatase inhibitors were not added to the buffer. 2xFLAG-Eps8 was eluted with wash buffer supplemented with 0.2 mg mL^−1^ 3xFLAG peptide and dialyzed into a buffer consisting of 25 mM MES pH 6.7, 75 mM NaCl, 5% glycerol and 1 mM DTT. The protein was concentrated and additionally purified using a Macro-Prep High S resin as above.

#### E. coli protein expression and purification

Human 14-3-3θ (UniProt: P27348) was expressed in *E. coli* BL21(DE3) cells using the IMPACT expression vector pTYB11 (New England BioLabs, Ipswich, MA), carrying a chitin-binding domain (CBD) for affinity purification and an intein domain for self-cleavage of the affinity tag after purification. Cells were grown in TB medium, supplemented with 100 mg mL^−1^ ampicillin at 37°C. Protein expression was induced with 1 mM IPTG at 18°C for 12 h. Cells were homogenized in 25 mM HEPES pH 7.5, 500 mM NaCl, 1 mM EDTA, 4 mM benzamidine hydrochloride and 1 mM PMSF, and lysed using a microfluidizer (Microfluidics, Newton, MA). Lysates were clarified by centrifugation at 20,000*g* for 30 min and the supernatant was loaded onto the chitin affinity resin (New England BioLabs). The affinity tag was removed by self-cleavage of the intein domain induced by incubation with 50 mM DTT overnight at room temperature. Proteins were eluted from the column in 25 mM HEPES pH 7.5, 50 mM NaCl, and 1 mM DTT. A final size exclusion chromatography purification step was performed on a Superdex 200 gel-filtration column (GE Healthcare, Little Chalfont, UK) in 25 mM HEPES pH 7.5, 50 mM NaCl and 1 mM DTT. The protein was concentrated to ~60 mg mL^−1^ using an ultracentrifugation filter (EMD Millipore, Burlington, MA). 14-3-3θ was also expressed as a maltose-binding protein (MBP) fusion in *E. coli* BL21(DE3). Cells were grown and lysed as described above, except cells were supplemented with 2 mg mL^−1^ glucose during growth. MBP-14-3-3θ was loaded onto an amylose resin, and the resin was washed with 10 column volumes of a buffer consisting of 25 mM HEPES pH 7.5, 300 mM NaCl and 1 mM DTT. MBP-14-3-3θ was eluted in 25 mM HEPES pH 7.5, 50 mM NaCl, 1 mM DTT and 10 mM maltose. The protein was additionally purified by size exclusion chromatography and concentrated as described above. The human Cdc42 (residues 1–178) constitutively active mutant G12V was also expressed in *E. coli* BL21(DE3) cells using the IMPACT expression vector pTYB11, as we have described ^10^.

### Synthetic IRSp53 phospho-peptides

Singly- and doubly-phosphorylated IRSp53 peptides (**Supplementary Table 1**) were synthesized with N-terminal acetylation and C-terminal amidation by Dr. Paul Leavis (Tufts University) and Neo Scientific (Cambridge, MA). Lyophilized peptides were dissolved at a concentration of ~10 mM and dialyzed extensively against 20 mM HEPES pH 7.5, 50 mM NaCl, 1 mM MgCl_2_, and 0.1 mM TCEP. The concentration of the peptides was determined using a fluorescence essay using fluorescamine-labeled peptide aliquots ^60^.

### Antibodies

Mouse monoclonal anti-IRSp53, rabbit polyclonal anti-pan-14-3-3 and mouse monoclonal anti-β-actin antibodies were purchased from Santa Cruz Biotechnology (Dallas, TX)

### Fluorescence

#### IRSp53 FRET sensor

We have previously described the overall design and data collection strategy of the IRSp53 FRET sensor expressed in *E. coli* cells ^10^. A critical modification here was the expression of the sensor protein (pIRSp53_FS_) in mammalian HEK293T cells, which together with limited starvation ensures proper phosphorylation at 14-3-3-binding sites (see above). Briefly, IRSp53 contains four endogenous cysteine residues, but only one (C230 within the BAR domain) is accessible to labeling with fluorescence probes, as determined by mass spectrometry. We introduced a second reactive cysteine near the C-terminus by site-directed mutagenesis (S519C mutant). To ensure that most of the donor probes are coupled to acceptor probes, we first under-labeled the protein with the donor fluorescein maleimide (Thermo Fisher Scientific), using a 0.9.1 molar ratio of probe to IRSp53 dimer for 30 min at 4°C. A two-fold molar excess of the acceptor rhodamine malemide (Thermo Fisher Scientific) was then added for 1 h. The same batch of probes was used for all the labeling reactions of this study. Unreacted excess probes were removed by two successive passages through Zeba Spin Desalting Columns (Thermo Fisher Scientific), equilibrated in 25 mM HEPES pH 7.5, 150 mM NaCl, 1 mM MgCl_2_, 5% glycerol and 1 mM DTT. The extent of labeling reaction was determined by comparing the protein concentration to the concentration of the dyes, estimated from the peak absorbance of the fluorophores, using as extinction coefficients ε(492 nm) = 83,000 cm^−1^ M^−1^ for fluorescein-5-maleimide and ε(542 nm) = 91,000 cm^−1^ M^−1^ for tetramethylrhodamine-5-maleimide. The IRSp53 dimer to total dye ratio was determined to be ~1:4, indicating that the two reactive cysteine residues on each subunit of the IRSp53 dimer were labeled. Steady-state fluorescence emission spectra were recorded at 20°C upon titration of GNPPNP-Cdc42 (G12V mutant), Eps8 and/or 14-3-3θ into a solution containing 0.02 pIRSp53_FS_ using a Cary Eclipse Fluorescence Spectrophotometer (Agilent Technologies). The experiments were performed in 25 mM Tris-HCl pH 7.5, 150 mM NaCl, 1 mM MgCl_2_, 5% glycerol and 0.1 mM TCEP. The excitation was set at 493 nm (10-nm slit width) and the emission spectra were recorded at 520 nm (10-nm slit width). Titration curves were fit using the IGOR Pro 7 data analysis software (WaveMetrics, www.wavemetrics.com) to either a non-cooperative binding function *F*([*ligand*]) = *F_max_* × [*ligand*] × (*K_d_* + [*ligand*])^−1^ or a cooperative binding function with non-specific binding *F*([*ligand*]) = *F_max_* × [*ligand*]*^n^* × (*K_d_^n^* + [*ligand*]*^n^*)*^−^*^1^ + *NS\**[*ligand*] (as indicated), where *n* is the Hill coefficient and *NS* is a constant non-specific fit parameter. Best fits were selected on the basis of residual analysis.

#### Peptide FRET

The Site 2 and Sites 2,4 peptides were fluorescently labeled at the amidated C-terminus. Both peptides were labeled with the donor probe Alexa Fluor 488 NHS Ester, whereas the Site 2 peptide was additionally labeled with the acceptor probe Alexa Fluor 568 NHS Ester (Thermo Fisher Scientific). The labeling reaction was performed at 25°C for 1 h in 20 mM MES pH 6.8 and 50 mM NaCl and at a peptide concentration of 5 mg mL^−1^ with a 2-fold molar excess of fluorescent dye. The labeling reaction was stopped with the addition of a 10-fold molar excess of hydroxylamine (Sigma-Aldrich Corporation). Labeled peptides were separated from any remaining unlabeled peptide using a Symmetry300 C_18_ reverse phase HPLC column (Waters Corporation, Milford, MA). The peptides were lyophilized, resuspended in 100 mM HEPES pH 7.5, and dialyzed against 25 mM HEPES pH 7.5, 50 mM NaCl and 1 mM MgCl_2_. The donor-labeled peptides were pre-incubated with 14-3-3θ at a 1:1 molar ratio *(i.e*. one peptide per 14-3-3θ dimer). The acceptor-labeled Site 2 peptide (at 5 μM concentration) was then titrated into the preformed complexes. The steady-state fluorescence emission spectra were recorded at 25°C, with an excitation wavelength of 495 nm (5-nm slit width) and an emission wavelength of 520 nm (5-nm slit width).

### Size exclusion chromatography–multi-angle light scattering (SEC–MALS)

14-3-3θ (100 μl at 4 mg mL^−1^) was pre-incubated for 2 h at 20°C with either the singly-phosphorylated Site 2 peptide at 1:2 molar ratio (one 14-3-3θ dimer per two Site 2 peptides) or the doubly-phosphorylated Sites 2,4 peptide at 1:1 molar ratio (one 14-3-3θ dimer per one Sites 2,4 peptide). The complexes were loaded onto a TSKgel SuperSW2000 column (Tosoh Corporation, Tokyo, Japan) connected in-line to a DAWN HELEOS MALS detector and an Optilab rEX (Wyatt Technology Corporation, Santa Barbara, CA) refractive index detector for mass analysis. Molecular masses were calculated with the program Astra (Wyatt Technology Corporation).

### Phosphoproteomics

Coomassie-stained gels were cut into 1 × 1 mm pieces, destained with 50% methanol and 1.25% acetic acid, reduced with 5 mM DTT, and alkylated with 20 mM iodoacetamide. The pieces were then washed with 20 mM ammonium bicarbonate and dehydrated with acetonitrile. Proteins were proteolyzed overnight at 37 °C with the addition of 5 ng mL^−1^ trypsin (Promega, Madison, WI) in 20 mM ammonium bicarbonate. Peptides were extracted with 0.3% trifluoroacetic acid, followed by a 50% acetonitrile/water solution. Tryptic peptides were separated by reverse phase HPLC chromatography using a 75 μm i.d. × 25 cm Acclaim PepMap nano LC column and analyzed by LC-MS/MS on a Q Exactive HF mass spectrometer (Thermo Fisher Scientific). The program Scaffold version 4.6.1 (Proteome Software, Portland, OR) was used for the identification of proteins and peptides, which were accepted when the identification probability was greater than 95%.

### Isothermal titration calorimetry (ITC)

ITC measurements were performed on a VP-ITC apparatus (MicroCal, Northampton, MA). Proteins and phospho-peptides were dialyzed side-by-side against 20 mM HEPES, pH 7.5, 150 mM NaCl, 1 mM MgCl_2_, and 0.1 mM TCEP (ITC buffer). The buffer was replaced three times over a period of two days of dialysis. Singly-phosphorylated peptides (at 100 μM) and doubly-phosphorylated peptides (at 50 μM) in the syringe were titrated into 14-3-3θ (at 10 μM) in the ITC cell of total volume 1.44 mL. All the experiments were carried out at 20°C. Titrations consisted of 10 μL injections, lasting for 20 s, with an interval of 5 min between injections. The heat of binding was corrected for the heat of injection, determined from either titrations of peptides into buffer (for the singly-phosphorylated peptides) or from the heat of injection at the end of the titration after reaching steady state (for the less abundant doubly-phosphorylated peptides). Each experiment was repeated three times. The data were analyzed using the program Origin (OriginLab, Northampton, MA). Best fits were selected on the basis of residual analysis. The integrated heats from all the binding reactions fit well to a single-site binding model.

### Crystallography

14-3-3θ was concentrated to 34 mg mL^−1^ in 20 mM HEPES pH 7.5, 50 mM NaCl, 1 mM MgCl_2_ and 1 mM DTT. Singly- and doubly-phosphorylated IRSp53 peptides were pre-incubated with 14-3-3θ at a molar ratio of 2.5:1 and 0.6:1, respectively, for 5 min at 25°C. Crystals were obtained at 18°C using the hanging-drop vapor-diffusion method. A typical 2μL hanging drop consisted of a 1:1 (v/v) mixture of the well solution and the 14-3θ/peptide complexes. For the crystallization of the singly-phosphorylated peptide complexes (Site 2, Site 3 and Site 4), the well solution consisted of 13-18% PEG 3350, 150 mM magnesium formate and 4% TFE. Crystals appeared after 1 day and were flash-frozen after a brief transfer into a cryo-solution containing 11 % PEG 400 and 11% PEG 1000 added to the crystallization buffer. For the crystallization of the doubly-phosphorylated IRSp53 peptide complexes (Sites 2,3 and Sites 3,4), the well solution consisted of 13-15% PEG 3350 and 100 mM CaCl_2_. Crystals grew over a period of 3 days and were slowly equilibrated into a cryo-solution containing 11% PEG 400 and 11% PEG 1000 over the course of 1 day. All the x-ray diffraction datasets were collected at 100 K and on three different sources (as indicated in **Table 1**): beamline 17-ID (APS, Argonne), beamline A1 (MacCHESS, Cornell University), and our home source (Bruker X8 Prospector, Billerica, MA). The datasets collected at synchrotron beamlines were indexed and scaled with the program HKL2000 (HKL Research) and the datasets collected at our home source were processed using the Bruker program SAINT (v8.34A). The structures were determined by molecular replacement, using as a search model PDB entry 2BTP (excluding from this model the bound peptide). Model building and refinement were performed with the programs Coot ^70^ and Phenix ^71^, respectively. Figures of the structure were prepared with program PyMOL (http://www.pymol.org/).

### Membrane fractionation

HEK293T cells grown in 15-cm plates were transfected with either IRSp53-FLAG or M234-FLAG. After 48 h expression, cells were incubated for 2 h in fresh growth media, which either contained or lacked 10% serum. Cells were then washed and incubated in PBS supplemented with 0.1 Calyculin A and 1x Halt Phosphatase Inhibitor Cocktail for 15 min on ice. Each cell plate was harvested, pelleted and resuspended in 250 fractionation buffer consisting of 20 mM HEPES-KOH pH 7.4, 10 mM KCl, 2 mM MgCl_2_, 1 mM EDTA, 1 mM EGTA, 1 mM DTT, 1 mM PMSF, 0.1 μM Calyculin A, 1x cOmplete protease inhibitor cocktail and 1x Halt Phosphatase Inhibitor Cocktail, before passing through a 27-gauge needle 10 times. Ruptured cells were incubated on ice for 20 min and the nuclei were removed by centrifugation at 720g for 5 min. Mitochondria were removed from the supernatant by centrifugation at 10,000*g* for 5 min, and the resulting supernatant was separated into cytosolic and membrane fractions by ultracentrifugation at 100,000*g* for 1 h. The cytosolic fraction was transferred to a fresh tube, and the membrane fraction was resuspended in fractionation buffer by passing several times through a 25-gauge needle. The membrane fraction was pelleted and resuspended two more times before resuspending the final membrane pellet in 50 μL TBS supplemented with 0.1% SDS, 1 mM PMSF, 0.1 μM Calyculin A, 1x cOmplete protease inhibitor cocktail and 1x Halt Phosphatase Inhibitor Cocktail. The relative abundance of IRSp53 was determined by densitometry analysis using the program Fiji performed of Western blots that were transferred from SDS-acrylamide gels, loaded with 2 μL of cytosolic fraction and 1.3 μL of membrane fraction for each condition. The fractional abundance of IRSp53 accounts for the difference in the concentration of cytosolic and membrane samples (which were resuspended in different volumes), the amount loaded onto the SDS-acrylamide gels, and the abundance of cytosolic GAPDH and membrane bound β-actin. All the experiments were performed in triplicate from three independent transfections.

### Statistical methods

The statistical significance of the measurements was determined using the program SigmaPlot 13 (Systat Software), using the unpaired two-sided Student’s t-test or Mann-Whitney’s rank sum test (as indicated). A p-value ≤ 0.05 was considered significant.

## Data availability

The coordinates and structure factors of the five structures of complexes of 14-3-3θ with singly- and doubly-phosphorylated IRSp53 peptides have been deposited in the Protein Data Bank (PDB) with accession codes 6BCR (Site 2), 6BCY (Site 3), 6BD1 (Site 4), 6BQT (Sites 2,3) and 6BD2 (Sites 2,4). Protein constructs and other data reported in this study are available from the corresponding author upon request

## Acknowledgments

This research was supported by National Institutes of Health grant R01 MH087950 to R.D. D.J.K. was in part supported by American Cancer Society grant PF-13-033-01-DMC. We thank Paul Leavis (Tufts University) for the synthesis of some of the peptides used in this study. X-ray data collection at beamline X6A of the National Synchrotron Light Source was supported by NIH grant GM-0080 and DOE contract DE-AC02–98CH10886. X-ray data collection at MacCHESS beamline A1 was supported by NSF grant DMR-1332208 and NIH grant GM-103485.

## Author Contributions

D.R.K. and R.D. participated equally in the experimental design and the preparation of the manuscript and figures. D.K. conducted most of the experiments, and R.D. participated in the determination of the crystal structures.

## Additional information

### Supplementary information

Supplementary Figures 1–4 and Supplementary Tables 1 and 2.

